# The PRALINE database: Protein and Rna humAn singLe nucleotIde variaNts in condEnsates

**DOI:** 10.1101/2022.12.03.518982

**Authors:** Andrea Vandelli, Magdalena Arnal Segura, Michele Monti, Jonathan Fiorentino, Laura Broglia, Alessio Colantoni, Natalia Sanchez de Groot, Marc Torrent Burgas, Alexandros Armaos, Gian Gaetano Tartaglia

**Affiliations:** Department of Biochemistry and Molecular Biology, Universitat Autònoma de Barcelona, Barcelona, 08193, Spain; Universitat Pompeu Fabra (UPF), 08003 Barcelona, Spain; Center for Human Technologies (CHT) and Center for Life Nano Science (CNLS), Istituto Italiano di Tecnologia (IIT), Via Enrico Melen, 83, 16152 Genova GE; Department of Biology and Biotechnologies, University Sapienza Rome, Via Aldo Moro 5, 00185, Roma, Italy

## Abstract

**Summary:** Biological condensates are membraneless organelles with different material properties. Proteins and RNAs are the main components, but most of their interactions are still unknown. Here we introduce PRALINE, a database for the interrogation of proteins and RNAs contained in stress-granules, processing bodies, and other assemblies including droplets and amyloids. PRALINE provides information about the predicted and experimentally validated protein-protein, protein-RNA and RNA-RNA interactions. For proteins, it reports the liquid-liquid phase separation and liquid-solid phase separation propensities. For RNAs, it provides information on predicted secondary structure content. PRALINE shows detailed information on human single-nucleotide variants, their clinical significance and presence in protein and RNA binding sites, and how they can affect condensates’ physical properties.

**Availability:** PRALINE is freely accessible on the web at http://alvinlee.bio.uniroma1.it/praline.

**Supplementary information:** General information is at http://alvinlee.bio.uniroma1.it/praline/about, where we provide a detailed description of the datasets and the tools employed in the database. Data provided in PRALINE are available at http://alvinlee.bio.uniroma1.it/praline/downloads. The tutorial is at http://alvinlee.bio.uniroma1.it/praline/tutorial.

## INTRODUCTION

Although the exact composition and functions of the different condensates are unknown, they are enriched in protein and RNA molecules that interact through protein-protein, protein-RNA and RNA-RNA networks. Solid-like condensates, and in particular amyloids, are generally considered to be inherently irreversible aberrant clumps (Dobson, 2017), while liquid-like condensates are dynamic entities that exchange components with the surrounding environment and grow, collapse and fuse in the nucleus and cytoplasm (Marchese et al., 2016). Liquid-like condensates perform different functions on RNA molecules such as storage in the germline, localization in neurons and protection from harmful conditions. The most known liquid-like condensates are processing bodies (PBs) and stress granules (SGs), both enriched in RNA that allows them to form and dissolve rapidly (Lorenzo Gotor et al., 2020). Yet, subtle changes in the composition or concentration of condensates’ constituents can induce the formation of solid-like assemblies (Cid-Samper et al., 2018). This is the case of Amyotrophic Lateral Sclerosis (ALS), where single-nucleotide variants (SNVs) in FUS trigger a liquid-to-solid phase transition (Patel et al., 2015). Structural properties of the RNA and changes upon mutations are important, since they play a role in the process of condensation. Highly structured RNAs attract large amounts of proteins thanks to their intrinsic ability to establish stable interactions (Sanchez de Groot et al., 2019). Moreover, RNAs can act as scaffolding elements (Armaos et al., 2021): whereas a polypeptide of 100 amino acids can interact with one or two proteins, a chain of 100 nucleotides is able to bind to 5–20 proteins (Vandelli et al., 2022). Poorly structured transcripts also induce condensation, as they base-pair with other RNAs establishing a dense network of contacts (Treeck et al., 2018). All this information is gathered in PRALINE, a database that provides information on different condensates’ components, their interaction networks, and disease-related variants.

## METHODS

### RNA set

All RNAs sequences were retrieved from Ensembl version 105 (Cunningham et al., 2022). 1841 Stress Granule (SG) enriched RNAs were taken from Khong et al. (Khong et al., 2017) selecting the ones with log2(fold-change) >=1 and p-value <=0.01. 4852 Processing Body (PB) enriched RNAs are taken from Hubstenberger et al. (Hubstenberger et al., 2017), retrieving the ones with log2(fold-change) >=1 and q-value <= 0.01) for a total of 5614 unique genes upon removal of obsolete gene ids in the ensembl version 105. For each gene, the longest isoform was taken. The CROSS algorithm for secondary structure prediction of RNAs was launched on the 5614 RNA sequences with Globalscore parameters (Delli Ponti et al., 2017).

### Protein set

997 UniprotKB IDs were retrieved from different sources. Stress Granule (SG) proteins were retrieved from Markmiller et al. (Markmiller et al., 2018), Youn et al. (Youn et al., 2018) and Jain et al. (Jain et al., 2016) studies. From Jain work we retrieved 411 proteins that were either already known SG proteins or newly discovered through mass spec and IF. From Markmiller and colleagues, we retrieved 397 proteins either already known from previous experiments or found with APEX technique in hek293, NPC and IPSC cells, as well as proteins found to be stress / cell specific or independent. 60 SG proteins were retrieved from Youn’s work. We retrieved proteins with Non-negative matrix factorization (NMF) values = 9 or in case of a different NMF value, we collected proteins found to co-localise with G3BP1.

Processing Body (PB) proteins were retrieved from Hustemberger et al. Hubstenberger and Youn et al. (Youn et al., 2018). From the Hustemberger’s study, we collected 125 proteins found to be significantly enriched in PBs with p-value < 0.025 as reported in the paper. From Gingras and colleagues’ work, we retrieved 42 proteins either with NMF value =8 or that co-localize with DCP1A.

From Vendruscolo and Fuxreiter work we retrieved 280 droplet forming proteins and 68 amyloid forming ones (Vendruscolo and Fuxreiter, 2022).

### SNVs

DisGeNet (release 7.0) curated variant-disease associations (Piñero et al., 2020) and ClinVar variants (Landrum et al., 2014; Mj et al., 2018) with a review status higher than one (“practice guideline”, “reviewed by expert panel” and “criteria provided, multiple submitters, no conflicts”) were downloaded in May 2022. From these datasets, we retrieved disease-related single nucleotide changes (SNVs). 13857 SNVs from DisGeNet and 48671 SNVs from ClinVar fell in the coding region of RNAs enriched in SG and PB and their relative position in the transcripts was retrieved from the Ensembl Variation 105 (Cunningham et al., 2022) in Human Short Variants dataset, excluding insertions and deletions. The CROSS algorithm with Globalscore parameters was launched on RNA fragments of 51 nt with the SNV in the center to calculate the difference in secondary structure content between the reference and alternative RNA sequences (Delli Ponti et al., 2017). To avoid smaller-size fragments, SNVs falling at the beginning and at the end of the RNAs were removed. The reference and alternative CROSS secondary structure propensity profiles of the 51 nt RNA fragments were represented against each other in plots. The mean difference in secondary structure between reference and alternative sequence was computed on a window size of 11 nt upstream and downstream the SNV. Information on the expression quantitative trait loci (eQTL) and splicing quantitative trait loci (sQTL) of the SNVs was obtained from GTEx V8 (GTEx Consortium, 2020).

### RNA-RNA interactions

In May 2020, we retrieved human RNA-RNA interactions from the RISE database (version 1.0) focusing on the experimental interactions obtained with PARIS technique (Gong et al., 2018). For each RNA, binding site location were mapped to the longest transcript in Ensembl 105, using blastn algorithm (Cunningham et al., 2022). We retrieved a total of 25.232 RNA-RNA interactions with at least one interactor being an enriched RNA in SG or PB. In 934 of those interactions we found at least a SNV located inside a binding site.

### Protein-RNA interactions

We provide experimentally determined protein-RNA interactions validated through eCLIP experiments available in May 2022 from https://www.encodeproject.org/eclip/ (Van Nostrand et al., 2020) as well as catRAPID predictions available in RNAct (Lang et al., 2019). For the experimentally validated interactions, binding sites are displayed.

### Protein-Protein interactions

As binding sites related to protein-protein interactions are not physically available, we provide links to an external database. Human curated protein-protein interactions are linked to BioGRID database version 4.4 (Oughtred et al., 2021).

### LLPS and LSPT Propensities

We computed the propensity to undergo liquid-like and solid-like condensation for the set of 997 proteins detailed previously and for their 6152 natural variants (involving 632 of them), retrieved from UniProtKB (release 2022_01) (The UniProt Consortium, 2021). We considered 5949 single point mutations and 203 deletions. To quantify the extent to which each sequence is prone to undergo LSPT and LLPS we used the Zyggregator (Tartaglia et al., 2008) and catGRANULE algorithms (Bolognesi et al., 2016), which compute the liquid-solid and liquid-liquid propensities, respectively.

We note that LLPS and LSPT are promoted by both intrinsic and extrinsic contributions. By intrinsic contributions we mean physico-chemical properties of the polypeptide chain such as the hydrophobicity for Zyggregator and the structural disorder for catGRANULE. The extrinsic contributions relate to environmental factors, such as concentration, pH, ionic strength, but not only, for instance crowding agents are also to be included. Our methods predict intrinsic properties of the polypeptide chain and do not take into account the different extrinsic contributions at present.

For the proteins that do not have natural variants present in UniProtKB we only report the WT scores of Zyggregator and catGRANULE.

## INTERPRETATION AND USE OF THE DATABASE

PRALINE can be accessed using protein or RNA names provided as: Gene Name, Ensembl Gene / Transcript ID (https://www.ensembl.org/) and UniprotKB ID (https://www.uniprot.org/; **Figure 1A**).

- Searching for a specific protein, the user can retrieve information on the condensate state (Droplet/liquid-like or Amyloid/solid-like) and the organelle in which it has been found (SG/PB). The predicted Liquid-liquid phase separation (LLPS) and liquid-solid phase transition (LSPT) propensities and profiles of the wild-type sequence are provided, calculated with *cat*GRANULE (Bolognesi et al., 2016) and Zyggregator (Tartaglia et al., 2008) methods, respectively (>0.80 accuracy in predicting regions of the proteins involved in protein condensation; **Figure 1B**). Experimentally validated protein-protein interactions are available through links to BioGRID (https://thebiogrid.org/; **Methods**), while experimental and predicted protein-RNA interactions can be retrieved from RNAct (https://rnact.crg.eu/; **Methods**). Protein-RNA interactions are calculated using *cat*RAPID, an algorithm trained on NMR and X-ray structures (AUC of 0.77 on eCLIP interactions) (Lang et al., 2019). The number of SNVs is shown for the protein of interest and, for each SNV, it is possible to interrogate the amino acid position, the difference in LSPT and LLPS propensities compared to the reference (i.e., wild-type protein) and retrieve information related to disease (Landrum et al., 2014; Piñero et al., 2020). LSPT and LLPS scores and profiles are provided (**Figure 1C**; **Methods**).
- Searching for a specific RNA, the user can retrieve information on the condensate state (SG/PB), the RNA secondary structure content (table and profile predicted using CROSS, http://s.tartaglialab.com/page/cross_group), the experimentally validated RNA interactions (RISE database, http://rise.life.tsinghua.edu.cn/) and the predicted or experimentally validated protein interactions reported in RNAact (https://rnact.crg.eu/) for both the reference sequence and SNVs (Mj et al., 2018; Piñero et al., 2020)(**Figure 1D,E**; **Methods**). The RNA-RNA interactions table reports information on different binding partners, if the interactors belong to a condensate, binding sites location in the transcripts and related SNVs (**Figure 1F**; **Methods**). The SNV section reports the position in the transcript, the difference in secondary structure compared to the reference (a numerical value and a profile image are provided) (Delli Ponti et al., 2017), associated diseases and interactions with RNAs (RISE database) as well as proteins (eCLIP https://www.encodeproject.org/eclip/; **Methods**) that involve the SNV containing region (**Figure 1F**; **Methods**).

**Figure 1.**
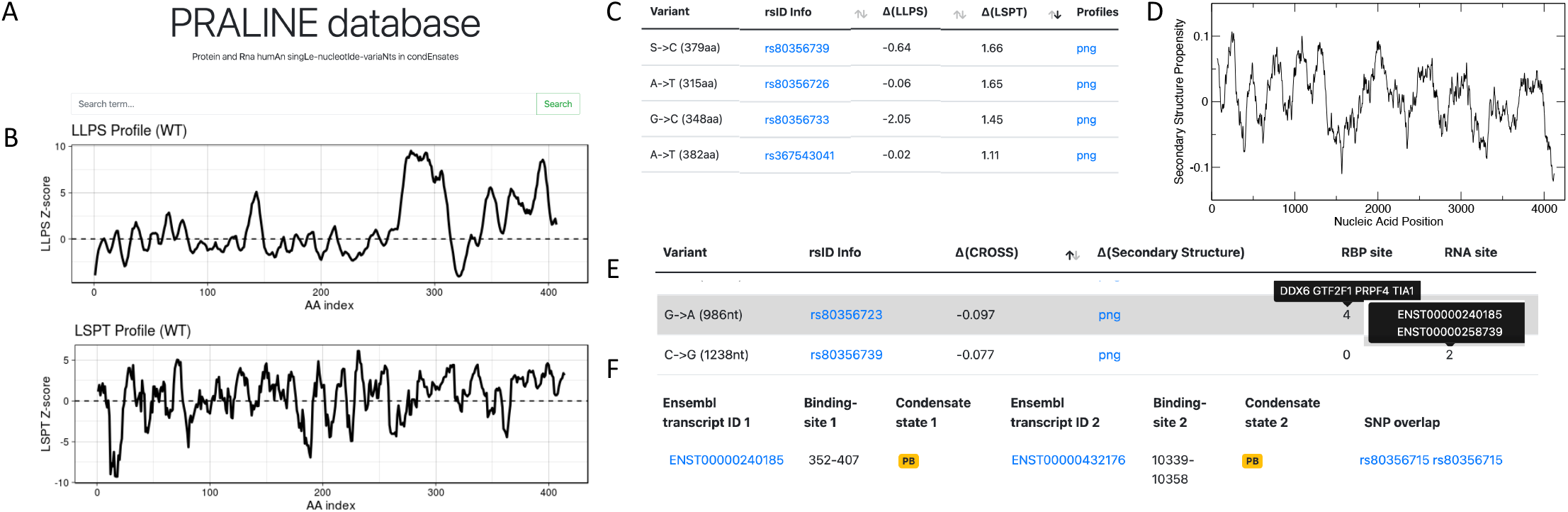
PRALINE database (**A**) Search bar. The input can be a protein or an RNA in different ID formats (**B**) Liquid-liquid phase-separation (LLPS) and liquid-solid phase-transition (LSPT) propensity profiles of a protein are predicted using *cat*GRANULE and Zyggregator algorithms. (**C**) Protein SNVs description table: the difference in LLPS and LSPT compared to the WT is provided. (**D**) CROSS secondary structure propensity profile image of a RNA sequence. (**E**) RNA SNVs description table: the difference in CROSS secondary structure propensity, compared to the WT, corresponding to a 11-nt window around the mutation is provided, as well as proteins and RNAs interacting with the query transcript containing the SNV. (**F**) Example of an RNA-RNA interaction table. The information about RNAs’ binding sites, condensates localization and SNVs falling inside at least one of the binding sites are reported. The examples **B-F** relate to *TARDBP*.

For most genes, information is available at both the protein and RNA levels, so it is possible to navigate from one molecule to the other, revealing the links between them.

## APPLICATIONS

*PRALINE* is a database that provides a comprehensive view of protein and RNA interactions and SNVs in human liquid-like and solid-like condensates. Information about experimentally validated and predicted molecular interactions, including protein-protein, protein-RNA and RNA-RNA, is provided, as well as the predicted RNA secondary structure content and both LLPS and LSPT propensities of proteins.

For each SNV, we provide a description of the associated diseases, the binding sites and the change in RNA secondary structure, LLPS and LSPT propensites. Combining physico-chemical properties of molecules and disease-related annotations, PRALINE helps to unravel macromolecular connections that sustain different types of condensates and how variants can affect their equilibrium. PRALINE is the first database providing LLPS and LSPT predictions for SNPs, and we envisage that it would greatly facilitate the design of experiments to study condensates’ formation and implication in human diseases. We note that although tested extensively and validated experimentally, *cat*GRANULE predictions could not be benchmarked against a database of individual SNVs causing LLPS, due to lack of adequate published resources. The availability of such databases will lead to a more precise understanding of the relationship between SNVs, structural conformations, protein-RNA assembly and diseases.

## ACKNOWLEDGEMENTS

The authors would like to thank Adriano Setti for the RNA-RNA interactions section and Leila Mansouri for the database name.

## Funding

Our research was supported by the ERC ASTRA_855923 and H2020 projects IASIS_727658 and INFORE_825080.

## Notes

http://alvinlee.bio.uniroma1.it/praline/about

### Competing Interest Statement

The authors have declared no competing interest.

