## Supplementary Material for "The PRALINE database: Protein and Rna humAn singLe nucleotIde variaNts in condEnsates"

### Tutorial

PRALINE database allows the study of proteins and RNAs found in assemblies associated with liquid-liquid phase separation and liquid-to-solid phase transition. PRALINE contains information about calculations and experimentally-validated macromolecular interactions as well as predictions of RNA secondary structure content, liquid-liquid phase-separation and liquid–to-solid phase transition propensities, with a focus on disease-related single-nucleotide variants (SNVs) contained in the molecules sequences.

PRALINE can be accessed using either protein or RNA names that can be provided as Gene Name, Ensembl Gene / Transcript ID or UniprotKB ID, as shown in the examples above.

PRALINE database

Protein and Rna humAn singLe-nucleotIde-variaNts in condEnsates

Search term... | Search

Search term examples:
RNA-binding protein (RBP) examples: FUS, TTR
RNA examples: ENST00000361789, ENST00000301030, ENSG00000035403
RBP/RNA example: ADAR, TARDBP

If a molecule (i.e TARDBP) is available both as a protein and as RNA, both types of information are shown, otherwise only one of them will be presented.

#### Proteins

| Ensembl gene ID | UniProtKB | Gene Name | Condensate state | Reference (PMID) | LLPS score | LSPT score | Protein - RNA binding sites (eCLIP\|RNAct) | Protein - Protein Interactions (BioGRID) | Variants |
|---|---|---|---|---|---|---|---|---|---|
| ENSG00000120948 | Q13148 | TARDBP | Droplet state (SG),Amyloid state | 29373831<br>26777405<br>34391803 | 1.64<br>(png\|txt) | 1.08<br>(png\|txt) | 6222(435) \| | | 26 |

#### RNAs

| Ensembl gene ID | Ensembl transcript ID | Gene Name | Condensate state | Reference (PMID) | Protein - RNA binding sites (eCLIP\|RNAct) | RNA - RNA Interactions | Double stranded Content | Variants |
|---|---|---|---|---|---|---|---|---|
| ENSG00000120948 | ENST00000240185 | TARDBP | Droplet state (PB) | 28965817 | 38(25) \| | 18 | 50.992%<br>(png\|txt) | 38 |

In the query page the user can find information about the condensate state in which the protein / RNA has been observed (Droplet, Amyloid), with a further SG/PB label if the molecule has been seen in stress granules (SGs) or processing bodies (PBs), respectively. The references of the studies reporting the molecule are shown as PMID. Indeed, by hovering over the column's name, each label is explained and relevant links to literature are provided.

#### Protein search

Searching for a protein, the user can retrieve the liquid-liquid phase separation (LLPS) and liquid-to-solid phase transition (LSPT) propensities, as predicted by catGRANULE and Zyggregator algorithms respectively. The scores are Z-normalised and displayed at single amino acid resolution (see About section for more information). Both profile graph and a table are available for the download.

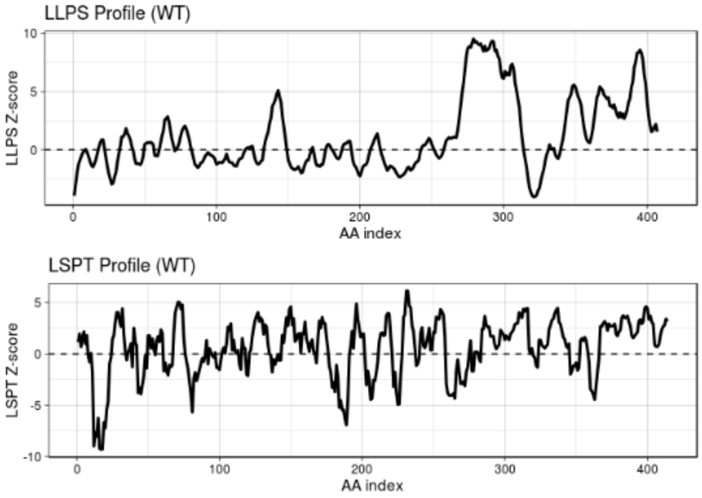


| aa_index | LLPS | LSPT |
|---|---|---|
| 1 | -3.93588688739327 | 1.15054052425111 |
| 2 | -2.844275 | 1.960643 |
| 3 | -1.8903091 | 0.6980005 |
| 4 | -1.215074 | 1.555846 |
| 5 | -0.8476039 | 2.1624051 |
| 6 | -0.5097959 | 0.9916056 |
| 7 | -0.2337287 | 1.7958671 |
| 8 | -0.06574037 | -0.19637125 |
| 9 | 0.03799738 | -0.79138435 |
| 10 | -0.24128937 | 0.05696414 |
| 11 | -0.6090509 | -1.821294 |
| 12 | -1.025915 | -8.987634 |
| 13 | -1.351346 | -7.654595 |
| 14 | -1.474272 | -8.025674 |
| 15 | -1.189684 | -6.259474 |
| 16 | -0.841708 | -9.224373 |
| 17 | -0.3254229 | -9.2910011 |
| 18 | 0.1880741 | -9.2910011 |
| 19 | 0.6052554 | -6.6930504 |
| 20 | 0.8636623 | -7.2218258 |

Information about experimentally validated protein-protein interactions is available through an external link to BioGRID database (33070389). BioGRID was not integrated in PRALINE because protein binding sites are not yet provided and the database is updated regularly.

The user can retrieve protein-RNA interactions predicted with catRAPID algorithm and available externally in the RNAct database, or experimentally validated interactions collected from eCLIP experiments (32728246). The user can visualize the total number of different interactions in which the protein is involved and between parentheses the number of contacts involving condensates RNAs.

#### TARDBP

Show 50 entries Search:

| RBP (Gene Name) | RBP Condensate state | Ensembl transcript ID | RNA Condensate state | Genomic Location | Cell line | FC (log) | p-value (log) | Annotation |
|---|---|---|---|---|---|---|---|---|
| TARDBP | Droplet state (SG),Amyloid state | ENST00000676644 | not in condensate | chr5:151806490-151806549 | K562 | 3.12896 | 9.85400 | exon |
| TARDBP | Droplet state (SG),Amyloid state | ENST00000677757 | Droplet state (PB) | chr5:151806490-151806549 | K562 | 3.12896 | 9.85400 | exon |
| TARDBP | Droplet state (SG),Amyloid state | ENST00000356245 | not in condensate | chr5:151806490-151806549 | K562 | 3.12896 | 9.85400 | exon |
| TARDBP | Droplet state (SG),Amyloid state | ENST00000677381 | not in condensate | chr5:151806490-151806549 | K562 | 3.12896 | 9.85400 | exon |

As in the other sections, by hovering over the column's name, the labels are explained and relevant links to literature are provided.

For each protein, a list of RNA partners is given, together with additional information, including the condensate state of both protein and RNA molecules, and chromosomal coordinates of the binding site and its annotation in the gene, as well as the cell line in which the analysis was done. Both the fold-change and P-value of the interactions are provided (27018577).

In the variants section, the SNVs (in rs format) collected from DisGeNet (31680165) and ClinVar (24234437) corresponding to the query protein (see About section for more information) are shown. For each SNV, the relative amino acid position inside the protein sequence is presented and the predicted difference in LLPS and LSPT of the mutant protein compared to the WT is calculated and shown in different plots.

## Q13148

Show 10 entries

| Variant | rsID Info | Δ(LLPS) | Δ(LSPT) | Profiles (LLPS and LSPT) |
|---|---|---|---|---|
| A->T (315aa) | rs80356726 | -0.06 | 1.65 | png |
| A->T (382aa) | rs367543041 | -0.02 | 1.11 | png |
| A->T (382aa) | rs367543041 | -0.02 | 1.11 | png |
| A->V (321aa) | VAR_083737 | 0.08 | 0.88 | png |
| A->V (90aa) | rs80356715 | 0.09 | 0.88 | png |
| A->V (90aa) | rs80356715 | 0.09 | 0.88 | png |
| D->G (169aa) | rs80356717 | 1.19 | -0.08 | png |
| G->A (290aa) | rs121908395 | -1.66 | -0.38 | png |
| G->A (290aa) | rs121908395 | -1.66 | -0.38 | png |
| G->A (294aa) | rs80356721 | -1.66 | -0.38 | png |

Showing 1 to 10 of 43 entries

Previous 1 2

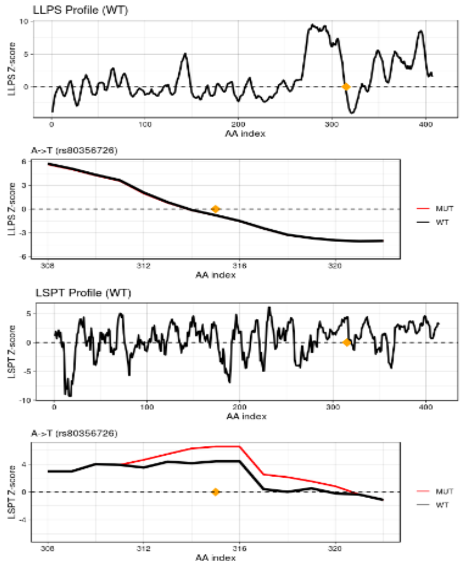

By clicking on a SNV name, the user can find additional information on the diseases related to the variant.

## rs80356726

##### Diseases

| rsID | Disease ID | Disease Name | Disease Class |
|---|---|---|---|
| rs80356726 | C2677565 | Amyotrophic lateral sclerosis 10 (disorder) | - |
| rs80356726 | | Amyotrophic lateral sclerosis type 10 | |
| rs80356726 | | Tardbp-related frontotemporal dementia | |

### RNA search

Searching for an RNA, the user can retrieve information about its condensate state and related references, as well as predicted and experimental protein-RNA interactions.

In addition, the predicted RNA secondary structure content obtained with CROSS algorithm is shown and a single-nucleotide propensity profile is available both as a graph and as a table. (see CROSS tutorial page for more information on the output files).

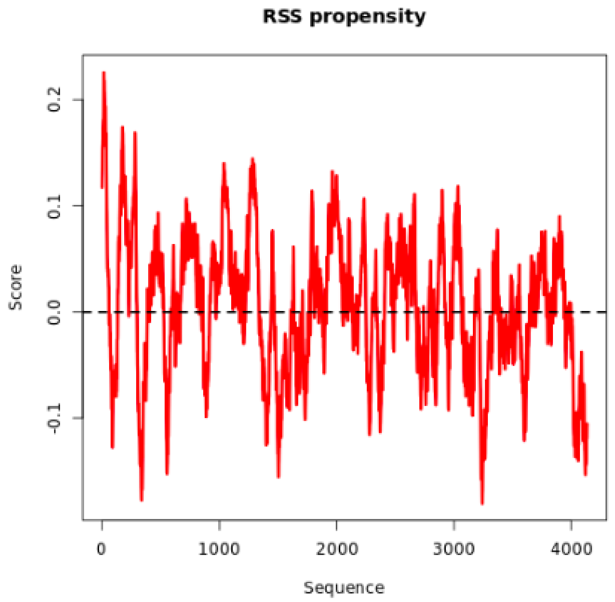


| | | | |
|---|---|---|---|
| 1 | A | -0.404 | - |
| 2 | T | -0.423 | - |
| 3 | T | -0.526 | - |
| 4 | T | 0 | - |
| 5 | T | 0.163 | - |
| 6 | G | -0.070 | - |
| 7 | T | 0.138 | - |
| 8 | G | 0.013 | - |
| 9 | G | -0.107 | - |
| 10 | G | -0.094 | - |
| 11 | A | -0.387 | - |
| 12 | G | 0.315 | - |
| 13 | C | 0.105 | - |
| 14 | G | -0.109 | - |
| 15 | A | -0.524 | - |
| 16 | A | -0.452 | - |
| 17 | G | -0.434 | - |
| 18 | C | 0.260 | - |
| 19 | G | 0.390 | - |
| 20 | G | 0.184 | - |
| 21 | T | 0.254 | - |
| 22 | G | 0.207 | - |
| 23 | G | 0.138 | - |
| 24 | C | 0.281 | - |
| 25 | T | 0.434 | - |
| 26 | G | 0.172 | - |
| 27 | G | 0.020 | 0.117262 |
| 28 | G | 0.373 | 0.1293035 |
| 29 | C | 0.416 | 0.1363238 |
| 30 | T | 0.391 | 0.1519443 |
| 31 | G | 0.228 | 0.1563469 |

High-throughput experimental RNA-RNA interactions retrieved from the RISE database are available. For each RNA the total number of interactors is shown together with the number of interactors belonging to condensates between parentheses. For each RNA pair, we show the condensate state information and the binding site location of both RNAs and the SNVs overlapping with at least one of the binding sites.

#### ENST00000240185

Show 10 entries

| Ensembl transcript ID 1 | Binding-site 1 | Condensate state 1 | Ensembl transcript ID 2 | Binding-site 2 | Condensate state 2 | SNP overlap |
|---|---|---|---|---|---|---|
| ENST00000240185 | 1735-1789 | PB | ENST00000260983 | 6865-6894 | SG-PB | NO_SNPs |
| ENST00000240185 | 2704-2725 | PB | ENST00000355086 | 13609-13640 | SG-PB | NO_SNPs |
| ENST00000240185 | 1341-1373 | PB | ENST00000562955 | 2289-2311 | SG | NO_SNPs |
| ENST00000240185 | 1727-1753 | PB | ENST00000297338 | 2906-2937 | PB | NO_SNPs |
| ENST00000240185 | 1727-1753 | PB | ENST00000297338 | 2906-2942 | PB | NO_SNPs |

As in the other sections, by hovering over the column's name, the labels are explained and relevant links to literature are provided.

In the variant section, we report the relative position of the SNV in the RNA, as well as the difference in secondary structure propensity of the mutant compared to the WT. The change in the propensity profile is also plotted. For each SNV, we report the RBPs and RNAs that interact with the query RNA in a region containing the SNV. As for the protein counterpart, clicking on the SNV name leads to more information about the diseases related to it.

##### ENST00000240185

Show 10 entries

| Variant | rsID Info | Δ(CROSS) | Δ(Secondary Structure) | RBP site | RNA site |
|---|---|---|---|---|---|
| A->G (1111nt) | rs80356730 | 0.062 | png | 1 | 0 |
| A->G (1130nt) | rs80356731 | 0.065 | png | 1 | 0 |
| A->G (1157nt) | rs80356734 | -0.037 | png | 0 | 0 |
| A->G (1249nt) | rs80356740 | 0.082 | png | 0 | 2 |
| A->G (1270nt) | rs80356741 | 0.000 | png | 0 | 2 |
| A->G (1271nt) | rs80356742 | 0.068 | png | 0 | 2 |
| A->G (608nt) | rs80356717 | -0.002 | png | 0 | 0 |
| A->G (777nt) | rs61741294 | -0.044 | png | 3 | 0 |
| A->G (889nt) | rs267607102 | -0.043 | png | 0 | 0 |
| A->G (902nt) | rs80356718 | 0.025 | png | 0 | 0 |

GTF2F1

Showing 1 to 10 of 60 entries

Previous 1 2 3 4 5 6 Next

##### ENST00000240185

Show 10 entries

| Variant | rsID Info | Δ(CROSS) | Δ(Secondary Structure) | RBP site | RNA site |
|---|---|---|---|---|---|
| A->G (1111nt) | rs80356730 | 0.062 | png | 1 | 0 |
| A->G (1130nt) | rs80356731 | 0.065 | png | 1 | 0 |
| A->G (1157nt) | rs80356734 | -0.037 | png | 0 | |
| A->G (1249nt) | rs80356740 | 0.082 | png | 0 | 2 |
| A->G (1270nt) | rs80356741 | 0.000 | png | 0 | 2 |
| A->G (1271nt) | rs80356742 | 0.068 | png | 0 | 2 |
| A->G (608nt) | rs80356717 | -0.002 | png | 0 | 0 |
| A->G (777nt) | rs61741294 | -0.044 | png | 3 | 0 |
| A->G (889nt) | rs267607102 | -0.043 | png | 0 | 0 |
| A->G (902nt) | rs80356718 | 0.025 | png | 0 | 0 |

ENST00000240185
ENST00000258739

Showing 1 to 10 of 60 entries

Previous 1 2 3 4 5 6 Next

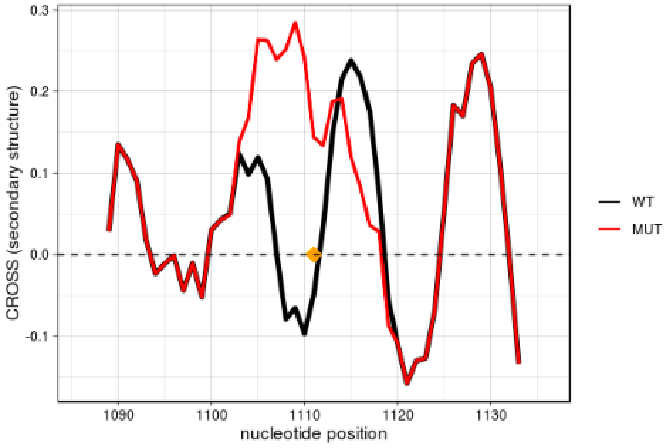


Finally in the Download section, PRALINE raw data for proteins, RNAs, their interactions and variants are available to the public.
